# Autopsy-based longitudinal multi-organ high-dimensional profiling reveals lineage plasticity in TRK-inhibitor-resistant secretory breast carcinoma

**DOI:** 10.64898/2026.04.06.716668

**Authors:** Yuki Muroyama, Mika Yanagaki, Hiroshi Tada, Akiko Ebata, Tateki Ito, Katsuhiko Ono, Jyunya Tominaga, Minoru Miyashita, Takashi Suzuki

## Abstract

Secretory breast carcinoma (SBC) is typically indolent, yet mechanisms underlying aggressiveness and therapeutic resistance to tropomyosin receptor kinase inhibitors (TRKi) remain unclear. Autopsy-based longitudinal multi-organ high-dimensional profiling of metastatic TRKi-resistant SBC demonstrated histopathological heterogeneity, including secretory and squamous components, arising from a shared clonal origin. Integrated genomic and transcriptomic analyses revealed hierarchical transcriptional rewiring consistent with a lineage-plastic state, suggesting a potential link to tumor aggressiveness and therapeutic resistance.

---

Secretory breast carcinoma (SBC) is a rare carcinoma characterized by *ETV6-NTRK3* gene fusion, accounting for less than 0.05% of invasive breast carcinomas^1^. SBC is typically indolent, and distant metastasis is rare^1^. Recently, tropomyosin receptor kinase inhibitors (TRKi), including larotrectinib and entrectinib, have demonstrated high objective response rates (57–80%) in NTRK fusion-positive solid tumors^2-4^. However, aggressive SBC with distant metastasis^5,6^, and TRKi resistance^7^ have been reported, and underlying molecular mechanisms remain poorly defined, representing an unmet clinical need.

Tumor heterogeneity contributes to therapeutic resistance across cancers, including clonal evolution and lineage plasticity^8^, the latter enabling transitions to alternative differentiation states under selective pressure, promoting phenotypic diversification and therapeutic resistance^9-13^.

However, clonal evolution and lineage plasticity across metastatic sites in aggressive SBC remain poorly characterized. Autopsy-based longitudinal multi-organ profiling enables controlled intra-patient analysis across metastatic sites, otherwise unachievable with conventional clinical sampling.

Here, we performed longitudinal multi-organ autopsy-based profiling of an aggressive TRKi-resistant SBC with marked histological heterogeneity, to elucidate clonal evolution and transcriptional reprogramming associated with lineage plasticity.

A 73-year-old woman (at autopsy, year X) underwent partial mastectomy with axillary dissection for right SBC at X−13 years with negative surgical margins (**Fig. 1A, Supplementary Fig. 1A**). Nine years later (X−4 years), a 6-cm right axillary lymph node mass was detected and confirmed as metastatic SBC on biopsy. The tumor was refractory to chemotherapy, and a right lung metastasis developed (**Fig. 1B, Supplementary Fig. 1B**). Comprehensive genomic profiling (CGP) using FoundationOne CDx confirmed the *ETV6-NTRK3* fusion in the axillary lesion. Microsatellite status was stable, and tumor mutational burden was low (6 mutations/Mb). Additional alterations included a *BRD4* P978_P980 deletion and a *TERT* promoter mutation (−124C>T). Entrectinib was initiated, resulting in a marked initial response, with reduction of axillary mass to 2 cm and lung metastasis regression (**Fig. 1B, Supplementary Fig. 1B**). However, the axillary lesion showed persistent fluorodeoxyglucose (FDG) uptake and subsequently progressed, with cystic change, necrosis, and ulceration extending to the skin (**Fig. 1B**). The lung metastasis recurred with multiple nodules (**Supplementary Fig. 1B**). Larotrectinib failed to control disease. The patient died in year X, and an autopsy was performed. Tissues from primary and metastatic sites were collected for histopathological and multi-omics analyses, including whole-exome sequencing (WES) and RNA sequencing (RNA-seq) (**Fig. 1A**).

**Fig. 1:**
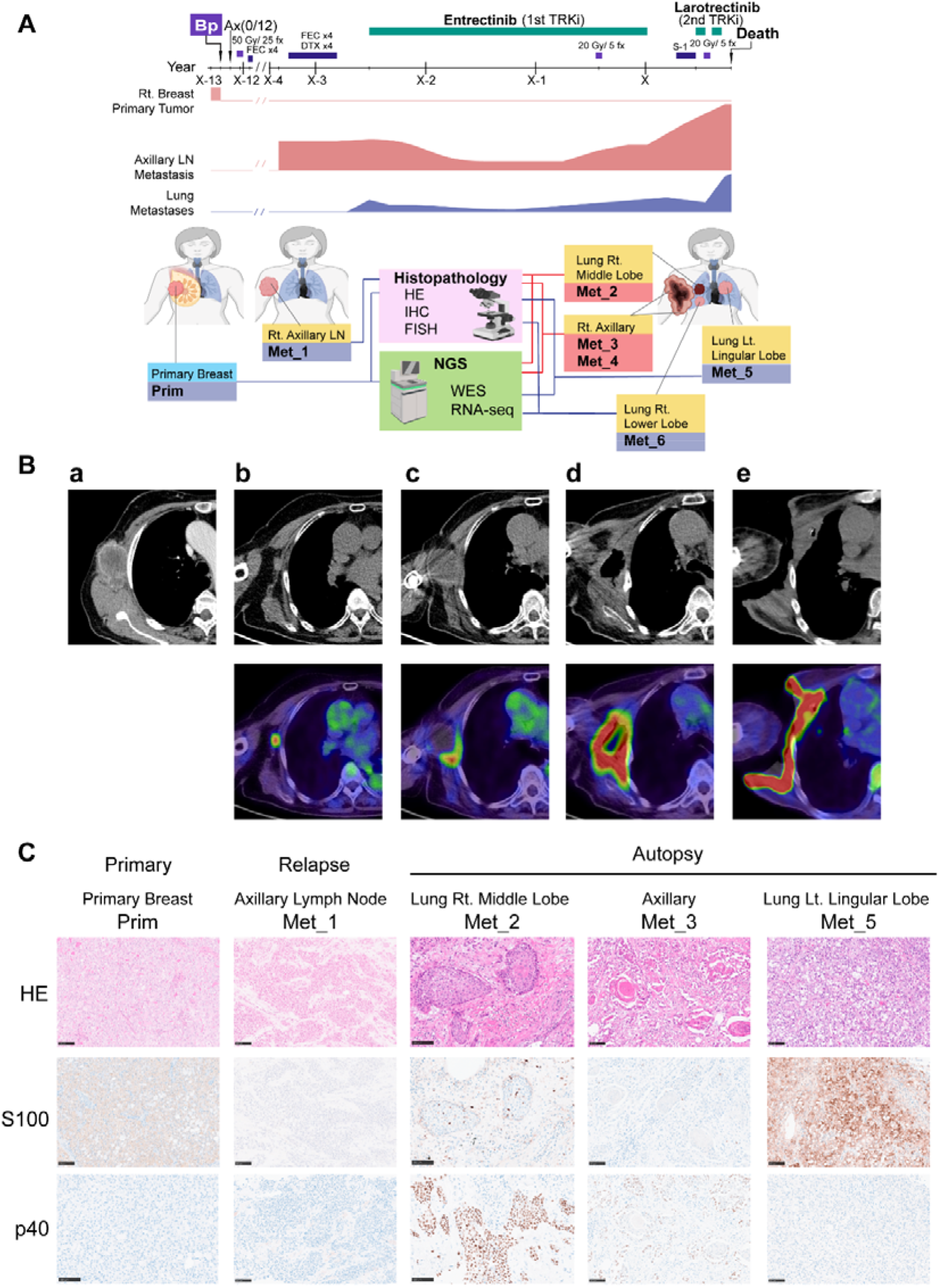
TRK inhibitor (TRKi)-resistant metastatic secretory breast carcinoma (SBC) exhibits histopathological heterogeneity. **(A)** Clinical timeline and sampling schema of a patient with metastatic SBC. Illustrations were created in part using BioRender.com. Rt: right, Lt: left, LN: lymph node, FEC: 5-Fluorouracil, Epirubicin and Cyclophosphamide, DTX: Docetaxel, fx: fraction. **(B)** Serial CT and FDG-PET/CT images demonstrating the temporal evolution of a right axillary lymph node metastasis (left to right). (a) Contrast-enhanced CT prior to TRKi therapy showing a large metastatic lymph node (approximately 6.5□cm) at X−3 years. (b)-(e) Corresponding FDG-PET/CT images are shown below each CT image. (b) Reduction in size (approximately 2□cm) with persistent FDG uptake after TRKi therapy (X−2 years). (c) Disease progression with cystic transformation (approximately 5□cm) at X−1 year. (d) Further progression with internal necrosis (approximately 10□cm) at X year. (e) Advanced progression with ulceration and extension to the skin surface (approximately 15□cm) at X year. **(C)** Representative histopathological features of the primary tumor and metastatic lesions. Hematoxylin and eosin (HE) staining and immunohistochemistry (IHC) for S100 (secretory marker) and p40 (squamous marker) are shown for the primary tumor (Prim), recurrent axillary lymph node (Met_1), right middle lobe lung metastasis (Met_2), axillary metastasis (Met_3), and left lingular lung metastasis (Met_5). Scale bars = 100□μm.

Histopathological analysis revealed spatiotemporal heterogeneity across metastatic sites (**Fig. 1C, Supplementary Fig. 2**). The primary breast tumor (Prim) showed vacuolated tumor cells arranged in microcystic patterns and was positive for S100, consistent with secretory carcinoma (Sec). The initial relapse axillary lesion (Met_1) retained Sec morphology but lacked S100 expression. At autopsy, axillary metastases (Met_3 and Met_4) and a right middle-lobe lung metastasis (Met_2) showed squamous differentiation with p40 positivity and loss of S100 staining, consistent with a squamous cell carcinoma (SCC)-like component. In contrast, the left lingular lung metastasis (Met_5) and right lower-lobe lung metastasis (Met_6) retained both histologic and immunohistochemical features consistent with Sec, demonstrating inter-metastatic heterogeneity. All regions were negative for ER, PgR, and HER2.

Histological diversity in TRKi-resistant SBC raised the question of whether the squamous components represented collision tumors or arose through clonal evolution. To address this question, we performed WES and phylogenetic tree construction, using Sequenza^14^, PyClone^15^ and MesKit^16^.

Phylogenetic reconstruction demonstrated that both secretory and squamous lesions share a common trunk (Public) origin shared by primary and metastatic tumors (**Fig. 2A**). Within the metastatic clade, the lesion retaining the secretory phenotype occupied a basal position, whereas squamous lesions occupied progressively more derived positions along the metastatic lineage, consistent with metastatic dissemination preceding squamous transdifferentiation.

**Fig. 2:**
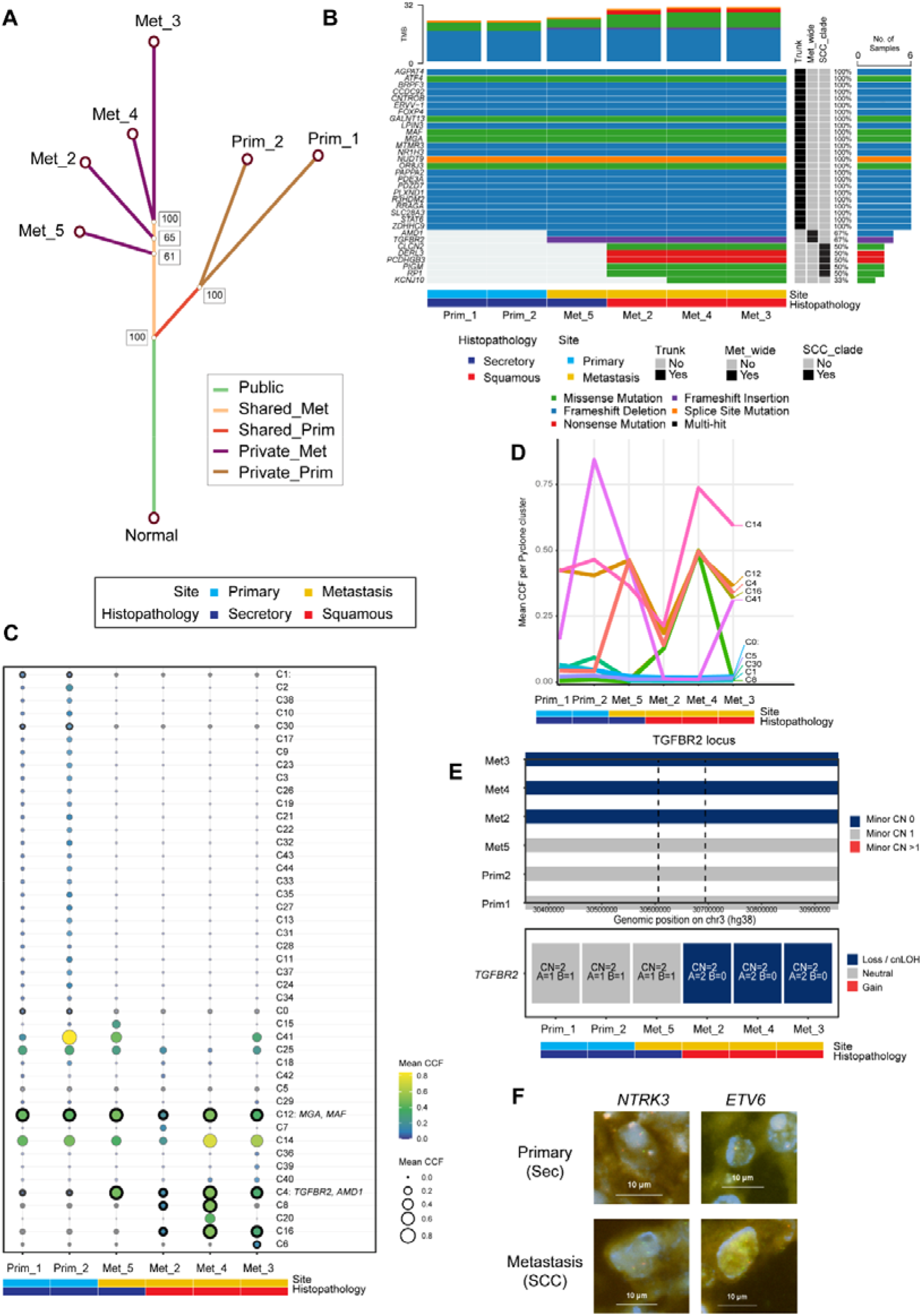
Genomic evidence for a shared clonal origin and lineage divergence between secretory and squamous components. **(A)** Phylogenetic reconstruction of primary and metastatic lesions using MesKit reveals a shared trunk (Public mutations) and distinct metastatic branches. Prim: primary, Met: metastasis. (Prim_1 and Prim_2 represent technical duplicates of the primary tumor.) **(B)** Oncoplot of high-impact somatic mutations showing trunk alterations present across all samples and metastatic-branch events, including metastasis-wide shared (Met_wide) and squamous-clade-restricted (SCC_clade) mutations. **(C)** Bubble plot of PyClone clusters across lesions ordered by the relative distribution of clones across samples. **(D)** Representative clonal trajectories across lesions, highlighting the expansion and persistence of major clones across samples. **(E)** Copy-number states at the *TGFBR2* locus across lesions, showing allele-specific alterations including copy-neutral loss of heterozygosity (cnLOH). CN: copy number, A: A allele, B: B allele. **(F)** Break-apart fluorescence in situ hybridization (FISH) demonstrating *ETV6* and *NTRK3* rearrangements in primary (secretory) and metastatic (squamous) lesions. Scale bars = 10 μm.

To identify genetic alterations underlying tumor evolution, we curated phylogenetically supported, high-confidence functional mutations (Methods) (**Fig. 2B**). These comprised 32 alterations, with 24 truncal events shared across all lesions and 8 metastasis-associated events, of which *TGFBR2* and *AMD1* alterations were shared across all metastases, whereas the remaining six were confined to squamous lesions.

To resolve subclonal dynamics, we analyzed PyClone-defined clusters as proxies for tumor subclones (**Fig. 2C, D**). Clusters were ordered based on their relative distributions across lesions. Clusters near the top of the plot were enriched in primary lesions, whereas more centrally positioned clusters, such as C12, harboring *MAF* and *MGA* gene alterations, were broadly distributed at high abundance across both primary and metastatic lesions, consistent with shared ancestral clones. In contrast, clusters such as C4, harboring *TGFBR2* alterations, were enriched in metastases, whereas later clusters, including C16, were preferentially expanded in squamous metastases. These patterns indicate stepwise subclonal diversification during metastatic progression.

Copy-number analysis revealed increased genomic complexity in metastatic lesions, particularly SCC lesions, characterized by extensive loss-of-heterozygosity (LOH) and copy-number alterations (CNA) (**Supplementary Fig. 3**). Allele-specific analysis identified copy-neutral LOH (cnLOH) at the *TGFBR2* locus (3p24.1) in squamous but not secretory lesions (**Fig. 2E**), indicating stepwise *TGFBR2* alterations, with metastasis-wide frameshift mutations followed by cnLOH in squamous metastases.

Canonical TRK inhibitor resistance-associated mutations, including secondary mutations in NTRK kinase domains or canonical bypass pathway alterations^17^, were not identified by WES.

Break-apart FISH confirmed *ETV6-NTRK3* gene rearrangements across secretory and squamous lesions (**Fig. 2F**), further supporting a shared clonal origin.

Together, these findings indicate that the histopathological heterogeneity observed in the TRKi-resistant SBC arises through clonal diversification from a common ancestor.

Next, we performed RNA-seq to dissect the transcriptional reprogramming between secretory and squamous states.

Principal component analysis (PCA) demonstrated clear segregation between Sec and SCC lesions along PC1 (73%), while PC2 (11%) separated primary and metastatic sites (**Fig. 3A**). Differential expression analysis identified genes upregulated in SCC, including *KRT17*, supporting squamous differentiation (**Fig. 3B, Supplementary Fig. 4**). Consistent with metastasis-wide frameshift indels and SCC-specific cnLOH, *TGFBR2* expression was reduced in SCC compared with Sec (**Fig. 3B, C**).

**Fig. 3:**
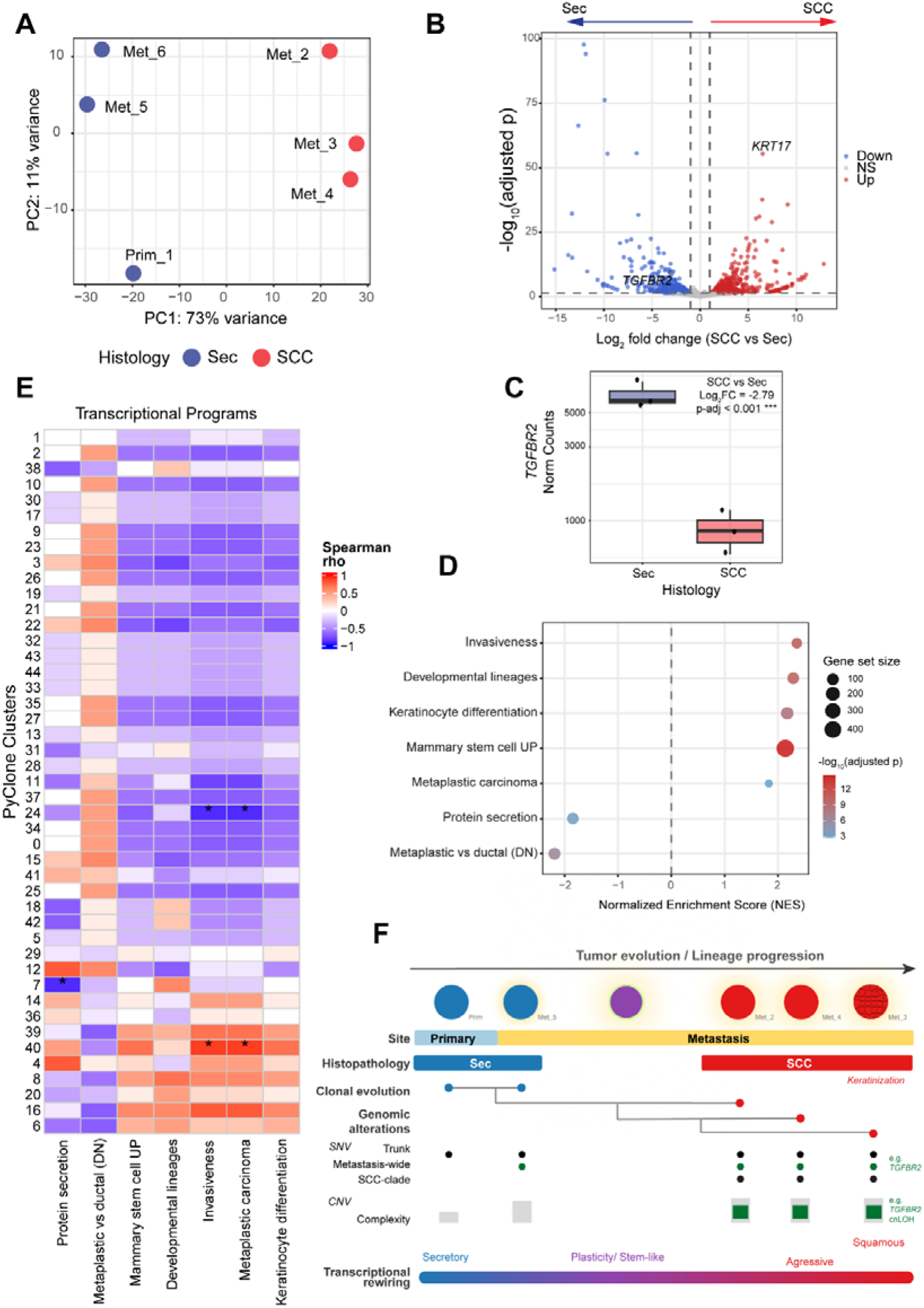
Transcriptional reprogramming associated with squamous differentiation. **(A)** Principal component analysis (PCA) of RNA-seq expression profiles from secretory carcinoma (Sec, n = 3) and squamous cell carcinoma (SCC, n = 3). **(B)** Volcano plot showing differential gene expression between SCC and Sec samples. Differential expression was estimated using DESeq2, and significance was assessed with Benjamini-Hochberg false discovery rate (FDR) correction. The y-axis represents −log_10_(adjusted P values). NS: non-significant. **(C)** Box plot showing *TGFBR2* expression levels (normalized counts) in secretory and squamous lesions (Sec, n = 3; SCC, n = 3). FC: fold change. Norm: normalized. **(D)** Gene set enrichment analysis (GSEA) of selected transcriptional programs using a pre-ranked gene list based on log2 fold changes (SCC vs Sec). Dot plots show normalized enrichment scores (NES), with dot size representing gene set size and color indicating −log_10_(FDR). FDR: false discovery rate. **(E)** Integration of clonal architecture and transcriptional programs. **(F)** Integrated model of tumor evolution and lineage plasticity in TRKi-resistant secretory breast carcinoma. SNV: single nucleotide variant. CNV: copy number variation

To delineate biological mechanisms underlying lineage transition, we performed gene set enrichment analysis (GSEA) (**Fig. 3D, Supplementary Table 1**). SCC lesions were enriched for squamous and keratinocyte differentiation and metaplastic breast carcinoma signatures, consistent with histology. They were also enriched for invasiveness signatures, as well as stem cell and developmental lineage signatures, indicating acquisition of lineage plasticity. Next, we integrated clonal evolution with transcriptional phenotypes across samples (**Fig. 3E**). Clusters broadly distributed across primary and metastatic lesions tended to be associated with transcriptional programs reflecting secretory epithelial function and non-metaplastic identity. In contrast, clusters enriched in metastatic and squamous lesions showed a tendency toward association with plasticity-related programs, including stem cell and developmental lineage signatures, alongside invasiveness and terminal keratinization programs. These trends indicate progressive transcriptional shifts along the clonal distribution gradient from secretory epithelial programs toward metaplastic phenotypes.

Taken together, the integration of genomic alterations, clonal architecture, and transcriptional pathway activity supports a model in which metastatic progression is accompanied by progressive transcriptional reprogramming toward lineage plasticity and squamous differentiation in TRKi-resistant SBC (**Fig. 3F**).

Leveraging autopsy-based multi-organ profiling, enabling controlled intra-patient comparisons across metastatic sites, we identified spatiotemporal heterogeneity in TRKi-resistant SBC. Phylogenetic reconstruction demonstrated that secretory and squamous components arise from a shared clonal origin, while integrated genomic and transcriptomic analyses revealed progressive transcriptional reprogramming toward metaplastic and squamous phenotypes, consistent with the acquisition of lineage plasticity.

Canonical TRKi-resistance-associated mutations, including those in kinase domains or bypass signaling pathways^17^, were not identified in this case, suggesting additional mechanisms of therapeutic resistance.

*TERT* promoter hotspot mutations, known to drive telomerase activation and tumor aggressiveness across multiple cancers^18^, have also been reported in aggressive SBC^5^. In our case, a *TERT* promoter mutation was identified in a metastatic lesion (Met_1) by CGP and may contribute to tumor aggressiveness but is unlikely to sufficiently explain the observed phenotypic diversification. Lineage plasticity has emerged as a key mechanism underlying phenotypic diversification and therapeutic resistance across cancers. Recent studies have highlighted that metastatic tumors can acquire plastic, fetal-progenitor-like status and differentiate into non-canonical states, such as squamous differentiation from adenocarcinoma, and that such plasticity is associated with tumor aggressiveness^13,19^. Our observations in the SBC case are conceptually consistent with these frameworks and collectively support a model in which lineage plasticity emerges progressively during metastatic evolution. Moreover, adeno-to-squamous transition has been shown to drive resistance to targeted therapy in lung cancer^12^. A recent case report also described squamous differentiation in TRKi-resistant SBC^7^, similarly to our case. Collectively, histological transdifferentiation toward a squamous phenotype may represent an adaptive process associated with aggressive disease progression and therapeutic resistance.

Among metastasis-associated alterations, *TGFBR2* frameshift insertions were observed across metastases, followed by cnLOH in squamous lesions with reduced transcript levels. Previous studies have implicated TGFBR2 signaling in epithelial plasticity, aggressive breast cancer phenotypes, and squamous carcinogenesis^20-23^, supporting a potential contribution to metastatic progression and lineage diversification in aggressive SBC.

Together, our findings suggest that lineage plasticity plays an underappreciated role in therapeutic resistance in SBC. Given the rarity of SBC and the distinctive aggressive clinical features, further accumulation of similar cases is warranted. Nevertheless, our autopsy-based profiling provides a framework for understanding adaptive evolution in rare fusion-driven tumors under targeted therapy.

## Abbreviations

SBC: secretory breast carcinoma
TRK: tropomyosin receptor kinase
NTRK3: Neurotrophic Tyrosine Kinase Receptor Type 3
ETV6: ETS Variant Transcription Factor 6
FEC: 5-Fluorouracil, Epirubicin and Cyclophosphamide
DTX: Docetaxel
fx: fraction
CGP: Comprehensive genomic profiling
Mb: megabase
CT: computed tomography
FDG: fluorodeoxyglucose
PET: positron emission tomography
IHC: immunohistochemistry
FFPE: formalin-fixed, paraffin-embedded
Sec: secretory carcinoma
SCC: squamous cell carcinoma
Rt: right
Lt: left
LN: lymph node
Prim: primary
Met: metastasis
WES: whole-exome sequencing
RNA-seq: RNA sequencing
CCF: cancer cell fraction
ER: Estrogen receptor
PgR: Progesterone receptor
HER2: Human epidermal growth factor receptor 2
S100: S100 calcium-binding protein
p40: ΔNp63 isoform of tumor protein p63 (TP63)
TGFBR2: Transforming Growth Factor Beta Receptor 2
AMD1: Adenosylmethionine Decarboxylase 1
MGA: Max-gene associated
SNV: single nucleotide variant
CNV: copy number variation
CNA: copy number alteration
cnLOH: copy neutral loss of heterozygosity
MAF: mutation annotation format
VAF: variant allele frequency
PCA: principal component analysis
DEG: differentially expressed gene
GSEA: gene set enrichment analysis
FDR: false discovery rate
NES: Normalized enrichment scores
NS: non-significant
DN: down

## Supplementary figure legends

**Supplementary Fig. 1: Radiological features of the primary tumor and longitudinal lung lesions**.

**Supplementary Fig. 2: Histopathological features of the primary tumor and metastatic lesions**.

**Supplementary Fig. 3: Genome-wide copy-number alterations**

**Supplementary Fig. 4: Heatmap of the top 50 differentially expressed genes (DEGs) distinguishing secretory and squamous lesions**.

**Supplementary Table legends:**

**Supplementary Table 1: Summary of gene set enrichment analysis (GSEA) for selected transcriptional programs**.

## Methods

Detailed methods are provided in the Supplementary Information.

## Ethical statement

Written informed consent was obtained from the patient’s next of kin. This study was approved by the Tohoku University Institutional Review Board (Protocol Identification No. 2025-1-649).

## Data availability

The datasets generated during and/or analyzed during the current study are available from the corresponding author on reasonable request.

## Author contributions

Y.M. conceptualized the study, wrote the original manuscript, prepared the figures, and contributed to data analysis.

M.Y., H.T., A.E., and M.M. were involved in patient care and clinical investigation.

Y.M., T.I., and T.S. performed pathological diagnosis and analyses.

K.O. performed histopathological and molecular experiments.

J.T. performed radiological interpretation and contributed to imaging figures.

Y.M., H.T., and T.S. supervised the study.

All authors reviewed and approved the final manuscript.

## Acknowledgements

This study was funded by the Center for Diversity, Equity and Inclusion (TUMUG) at Tohoku University (Project to Promote Gender Equality and Female Researchers). The funder played no role in study design, data collection, analysis and interpretation of data, or the writing of this manuscript. We thank members of the Department of Pathology, Tohoku University Hospital, the Department of Anatomic Pathology, Tohoku University Graduate School of Medicine, and the SiRIUS Institute of Medical Research for their support during this study. We thank Dr. Yoshiaki Onodera (Department of Anatomic Pathology), Dr. Shiori Fujisawa (the SiRIUS Institute of Medical Research), Dr. Asumi Yamazaki (Department of Breast and Endocrine Surgical Oncology, Graduate School of Medicine, Tohoku University), and Dr. Keigo Murakami and Dr. Toru Furukawa (Department of Investigative Pathology, Tohoku University Graduate School of Medicine) for their assistance and support.

